# A spatial representation of information underlies probabilistic computation in cells

**DOI:** 10.1101/2025.02.12.637580

**Authors:** Matt Thomson, Zitong Jerry Wang

## Abstract

Seeking a signal source in unstructured environments is a fundamental challenge in robotics. Similarly, cells in tissues track signal sources using noisy, fragmented molecular gradients, shaped by fluid flow and extracellular matrix interactions. However, the precise algorithm cells use for source seeking is unknown. We show that cells can perform source seeking using a biophysical implementation of a computational algorithm called Bayes filtering. Specifically, the spatial distribution of molecules within the cell encodes a probability distribution over source location, and intracellular transport processes update this distribution. Live-cell imaging and spatial proteomics reveal that receptor dynamics in vivo matches the evolution of belief distributions under Bayes filtering. Unlike standard Bayes filtering, the cellular implementation adapts to fluctuating measurement noise without explicitly estimating noise statistics. When translated to traditional robotics algorithms, this cell-inspired adaptation enables robust navigation without continuously estimating signal statistics. Our results show that cells can leverage spatial organization to implement probabilistic algorithms, bridging cellular behavior and engineered systems.

---

Source seeking is classic robotics task, where a robot must localize toward the source of a signal. For example, aerial drones are used to find the source of gas leaks (*1*), or underwater robots are deployed to identify chemical spills. In practice, uncertain environmental conditions like turbulent flow can break continuous signal gradients into discontinuous patches (*2*), where deterministic algorithms like gradient descent risks trapping robots in local signal patches (Figure 1A).

**Figure 1:**
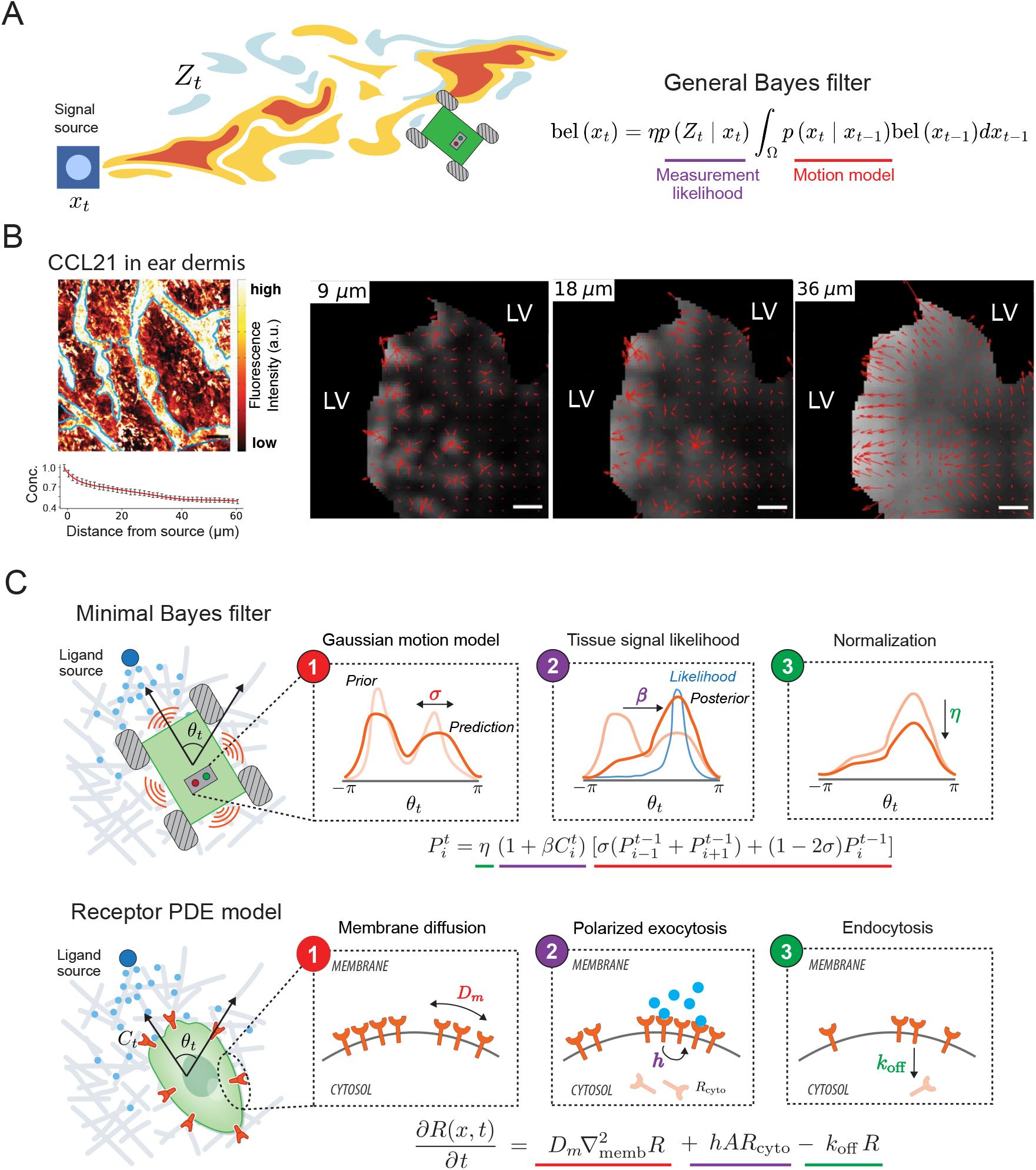
Mapping receptor dynamics to Bayes filtering. (**A**) General Bayes filter update equation for source seeking tasks such as plume tracking. (**B**) (Left) ECM-bound CCL21 in mouse ear dermis, and mean signal intensities relative to average maximum signal ± SEM as function of distance from the nearest LV margin. (Right) Red vectors indicate the direction and magnitude of local [CCL21] increase, computed by averaging over a circular surface area corresponding to different virtual cell size. Grayscale indicates the mean CCL21 intensity within each area. Scale bar, 15µ*m*. CCL21 images adapted from (*7*). (**C**) Mathematical equivalence between three steps of a Bayes filter update and three intracellular processes of a receptor PDE model.

Probabilistic methods for source seeking are often more effective than deterministic algorithms in unstructured environments, but are known to be computationally-intensive (*3*). In these algorithms, the agent maintains a probability distribution bel(*x*_*t*_) over all possible source positions *x*_*t*_ ∈ Ω and iteratively updates this belief using sensor measurements *Z*_*t*_ (Figure 1A). The core update rule underlying most such algorithm is given by the Bayes filter (*3*):

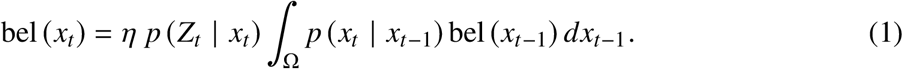

which yields the minimum mean squared error estimate of the true target position assuming accurate models. The algorithm consists of three steps: a prediction step to account for uncertainty in movement, where the prior belief bel (*x*_*t*−1_) is convolved with the motion model *p*(*x*_*t*_ | *x*_*t*−1_), followed by an update step to incorporate new sensor reading, where the belief is reweighted by the measurement likelihood *p*(*Z*_*t*_ | *x*_*t*_). Lastly, a normalization term *η* ensures belief sums to one. While effective, this iterative process is resource-intensive, requiring integration over the state space at each step and memory to store a full belief distribution over all candidate spatial locations.

Cells can efficiently localize to ligand sources in complex tissue environments, overcoming local signal peaks despite relying on noisy chemical hardware and operating under tight resource budgets. In tissues, extracellular matrix (ECM) binding and interstitial fluid flow break ligand gradients into irregular, fragmented patches (*4–8*). For example, CCL21, a chemokine secreted by lymphatic endothelial cells to guide cells toward lymphatic vessels, is transported by fluid flow and captured by a non-uniform ECM network. Quantitative imaging of mouse ear dermis shows that CCL21 forms a stable, reticulated pattern with local concentration peaks (*7*). CCL21 gradients appear smooth when averaged over tissue-scale regions, but vector field of local gradients, computed at the scale of individual cells (Figure 1B), reveal highly conflicting cues where local gradients often fail to align with the true source direction. These patchy gradients are stable over time due to strong ECM binding (*9*), having been observed for other morphogens and chemokines in vivo (*10–12*). This navigation problem is analogous to non-convex optimization, where naïve gradient descent only achieves local optima. Despite these challenges, live-cell imaging shows that cells can efficiently reach ligand sources (*7*), suggesting they use strategies that go beyond simple gradient-following.

Recent observations suggest dynamic spatial rearrangement of surface receptors may be important for source seeking tasks such as tracking and navigation. In neuronal growth cones, receptors such as Robo1 and PlxnA1 reorganize according to local ligand distributions, and inhibiting their rearrangement impairs directional guidance (*13,14*). In budding yeast, pheromone receptors dynamically rearrange to track the pheromone source during mating. In mesenchymal stem cells, blocking CCR2 receptor redistribution, without changing its overall expression, severely disrupts targeted migration to injured muscle tissues (*15*). These observations suggest that receptor dynamics, and not just expression, can be pivotal for robust source seeking.

In this work, we show that dynamic receptor rearrangement can function as a biophysical implementation of the Bayes filtering algorithm optimized for source seeking in complex environments. The spatial distribution of receptors encodes a probability distribution over source location and intracellular transport processes can update this distribution, thereby implementing key steps of the algorithm. Live-cell imaging and spatial proteomics data show that the Bayes filtering update rule accurately predicts the spatiotemporal dynamics of cell-surface receptors during source seeking. Unlike conventional Bayes filtering, the cellular implementation adapts to fluctuating measurement noise without explicitly estimating noise statistics. We show that translating this cell-inspired adaptation back into traditional robotics algorithms enables robust source seeking without the need to continuously estimate signal statistics. Our results illustrate how cells leverage spatial organization to implement complex, probabilistic algorithms.

## Receptor redistribution can implement Bayes filtering

We show that cells can implement an exact Bayes filter using known intracellular transport processes. To make this mapping concrete, we first construct a minimal Bayesian filter that is memory-efficient and computationally tractable (Figure 1B). We define the hidden state as the source direction *θ*_*t*_ ∈ [−*π, π*) relative to the agent’s location. This choice, rather than using the full coordinate space, minimizes memory demands. We discretize this variable into *N* possible directions, *θ*_*t*_ = 2*πi*/*N* for *i* = 1, ∼, *N*, with belief 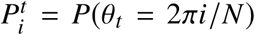 and input signal *C*_*t*_ ∈ ℤ^*N*^ representing ligand counts across membrane sectors. To simplify the prediction step, we use a Gaussian motion model to capture stochastic shifts in the source direction given cell movement:

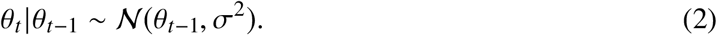

Assuming small variance reduces the integral in Equation (1) to a weighted sum:

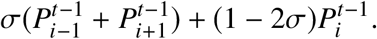

Next, we assume the measurement likelihood depends only on the ligand level in the source direction being conditioned on. We obtain a linear approximation of the likelihood model by fitting to signals sampled from interstitial gradients (*16*):

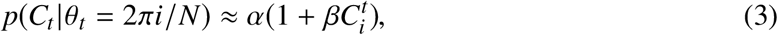

where 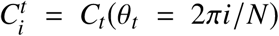. This model expresses how likely a ligand profile is, given the true source location. Note that *β* is small but positive, indicating that higher ligand counts only marginally increase the likelihood of a source, reflecting the patchiness of the signals. Substituting the motion model and measurement likelihood into Equation (1) yields a minimal Bayes filter for source seeking.:

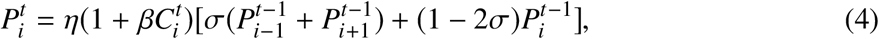

where *η* absorbed the constant factor from Equation (3) and represents the normalization step of the algorithm.

Receptor redistribution provides an exact biophysical implementation of the Bayesian filtering algorithm, where the spatial distribution of receptors encodes the belief distribution, and intracellular transport processes update it over time (Figure 1C). We show in the SI (*16*) that the minimal Bayes filter update of Equation (4) is mathematically equivalent to a standard partial differential equation (PDE) model describing receptor transport:

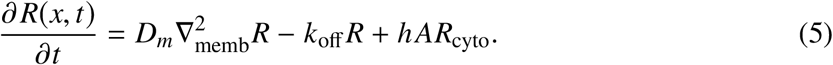

Here *R*(*x, t*) denotes the receptor concentration along the cell surface at location *x* and time *t*. This receptor profile represents the posterior belief *P* in the Bayes filter, as the rate of change of receptor across the cell surface *∂R*/*∂t* is equivalent to the Bayes belief update (Equation 4). Each step of this filtering algorithm maps to a term in the PDE. The motion model (Equation 2) maps to lateral receptor diffusion, represented by the Laplacian 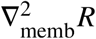, where membrane diffusivity *D*_*m*_ corresponds to the motion model variance *σ*^2^. The measurement likelihood (Equation 3) maps to exocytosis from a cytosolic pool at rate *h AR*_cyto_, polarized toward regions with high receptor activity creating a positive feedback. Here, likelihood model parameter *β* corresponds to rate constant *h*. Lastly, normalization (*η*) maps to receptor endocytosis with rate *k*_off_ (Figure 1C). In this way, filter updates of the belief correspond to receptor redistribution on the cell, and cells moving in the direction of maximal receptor activity approximately follows the maximum a posteriori estimate of the true source direction.

Polarized exocytosis of receptors for guidance cue (*14,17,18*), pheromones (*19*), chemokines (*15, 20*), and growth factors (*21, 22*) involve multiple intermediate processes, which could vary between receptors (*23*). For example, activation of DCC receptors by chemoattractant Netrin-1 lead to ligand-dependent clustering of DCC/Sytx1 complexes in activated membrane domains. Here, the formation of a SNARE complex between Sytx1 and TI-VAMP proteins occurs, thereby promoting exocytosis of DCC-containing vesicles at DCC-activated domains (*24*). For nerve growth factor (NGF), local activation of NGF receptors in growth cones induces asymmetric vesicle trafficking and targeted insertion of additional NGF receptors into the membrane near the site of stimulation via VAMP2-dependent exocytosis (*21*). Note some receptors may not redistribute in the manner described above (*25, 26*), since cells may implement Bayes filtering through membrane-bound effectors downstream of receptor activation rather than through receptor movement (see Discussion).

### In vivo receptor rearrangement matches evolution of Bayes belief distribution

Live-cell imaging and spatial proteomics show that Bayes filtering accurately predicts cell-surface receptor dynamics during source seeking (Figure 2A).

**Figure 2:**
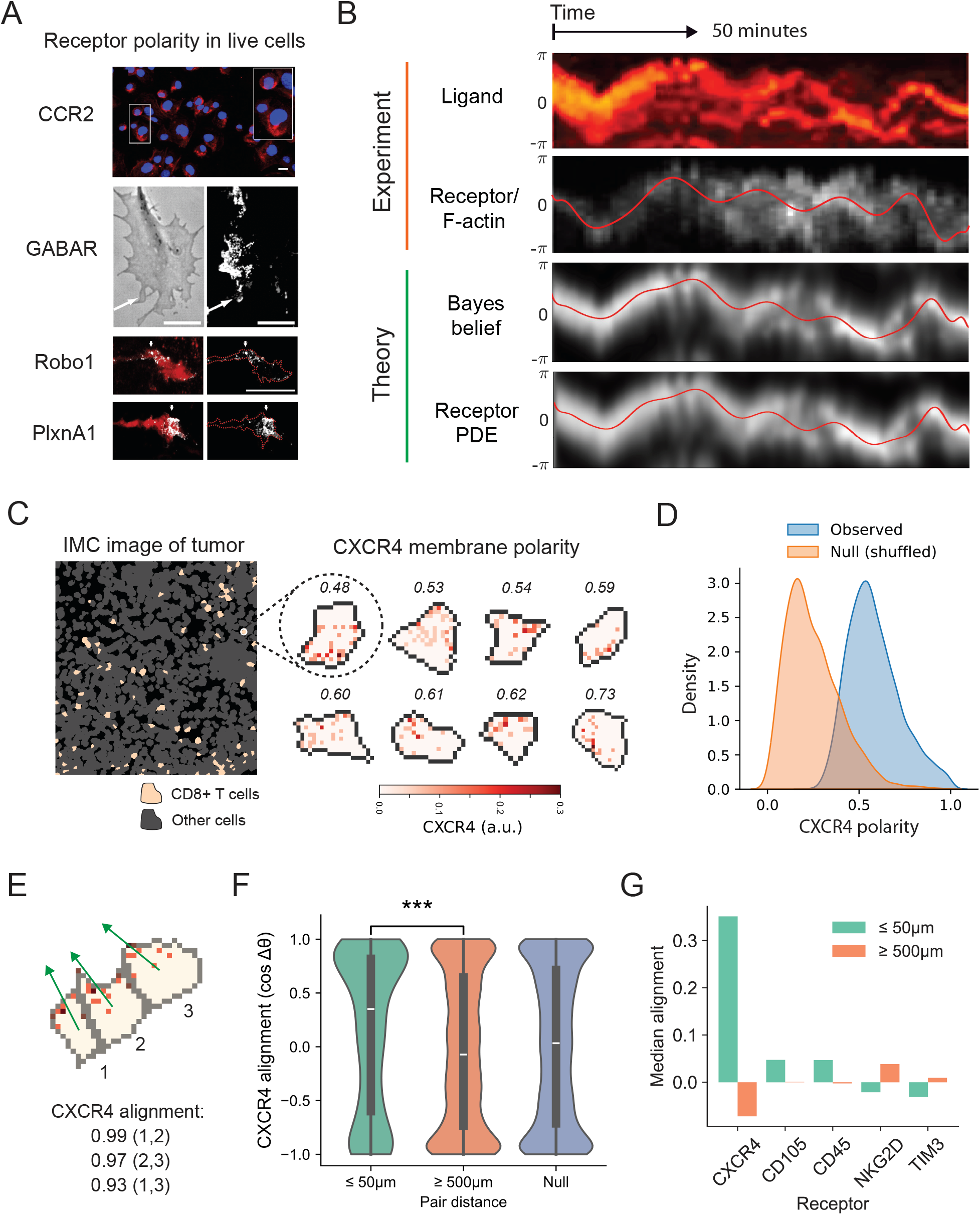
Bayes filtering update equation predicts in vivo receptor rearrangement. (**A**) Live-cell imaging of receptor membrane distribution, white arrow in GABAR image indicates direction of ligand gradient. Scale bar: 10µ*m*. Images adopted from (*14, 15, 18*). (**B**) Kymographs of UNC-6 present around the surface of an anchor cell during invasion, membrane distribution of f-actin in the same cell (highly correlated with UNC-40). Images adopted from (*27*). Belief distribution and receptor PDE output generated using ligand kymograph as input. (**C**) Example IMC image with example cells showing CXCR4 polarization. (**D**) Observed distribution of CXCR4 polarity among T cells in IMC dataset, Null distribution generated by randomly shuffling membrane pixels within individual cells. (**E**) Example of a T cell cluster with strongly aligned CXCR4 polarization found in IMC images. (**F**) Violin plots showing distribution of alignments computed between all pairs of T cells (cosine of difference in polarity angle) within 50µ*m* or greater than 500µ*m* apart. Null distribution represents randomly generated vector pairs. (**G**) Median alignment for different receptors.

In living cells, the spatial distribution of migratory receptors such as DCC evolves in a manner that mirrors Bayesian belief updates (Figure 2B). In animals, DCC (deleted in colorectal cancer) directs growth cone migration towards its extracellular ligand netrin. Live-cell imaging of the DCC orthologue UNC-40 in *C. elegans* shows that UNC-40 on the surface of anchor cells polarizes toward UNC-6 (netrin) during invasion and continuously redistributes in response to changing extracellular ligand distributions (Figure 2B) (*27*). Notably, when we feed the ligand kymograph as input to the Bayesian filtering algorithm, the evolution of the belief distribution (Equation 4) closely mirrors the observed receptor/F-actin distribution on the cell membrane (Figure 2B). This belief update also matches receptor dynamics simulated by the receptor PDE model (Equation 5), supporting the fact that cells implement Bayesian filtering through receptor rearrangement.

We find a similar pattern of receptor polarization in T cells from intact human tissue. Single-cell spatial proteomic profiling using Imaging Mass Cytometry (IMC) reveals that chemokine receptors, CXCR4, are significantly polarized in CD8+ T cells found in tumor (*28*) (Figure 2C). We quantified receptor polarity as the normalized vector sum of membrane intensity:

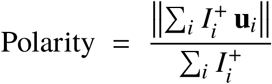

where 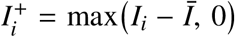 is the contrast-enhanced receptor intensity at the *i*-th membrane pixel, *Ī* is the mean membrane intensity, and **u**_*i*_ is the unit vector from the cell centroid to pixel *i*. For cells localizing to signal source, the Bayes filtering equation predicts that 1) signal receptor distributions should be polarized rather than uniform and 2) cells in similar local environments should exhibit aligned receptor polarity. As predicted, observed CXCR4 polarity in T cells is significantly higher than in randomly shuffled controls (Figure 2D). Furthermore, CXCR4 polarity vectors of nearby T cells (≤ 50µ*m*) are strongly aligned (p-value: 0.0009), whereas those of distant cells (≥ 500µ*m*) are randomly oriented (Figure 2E,F). This distance-dependent alignment is specific to the chemokine receptor: other surface proteins such as TIM3 and CD45 show no such pattern (Figure 2G), demonstrating that CXCR4 distributions are actively shaped by ligand distributions in vivo as predicted by the Bayes filtering equation.

### Receptor redistribution overcomes patchy gradients

Simulations show that Bayes filtering and its biophysical implementation via receptor redistribution enable robust navigation in patchy interstitial gradients.

To model these challenging environments (*16*), we simulate fluid flow carrying signaling ligands through an irregular ECM network, generating realistic patchy ligand distributions (Figure 3A). These simulated patterns closely resemble experimentally observed chemokine gradients (Figure 1B). In both tissue imaging and simulation, local gradient directions experienced by a typical cell (10-20 µm in diameter) fail to align with the global gradient direction, whereas larger cells (40µm) do not experience this misalignment (Figure 3B).

**Figure 3:**
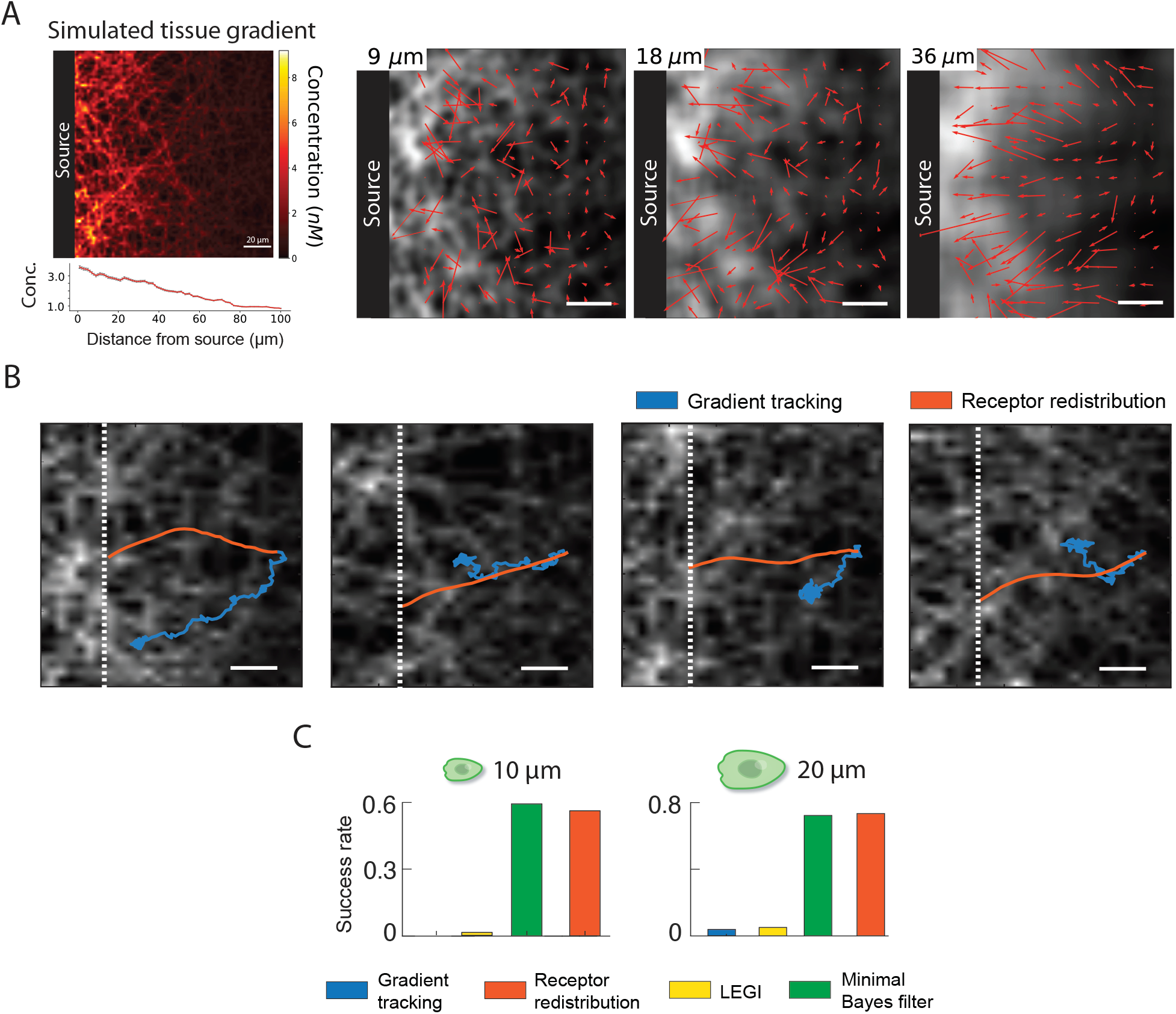
Comparing navigation strategies in patchy gradient. (**A**) Simulated ECM-bound gradient (right). Vector maps of local gradients as observed by virtual cells of a given diameter; Scale bar, 15µ*m*. (**B**) Simulated trajectories for cells following local gradient vs. using receptor redistribution; Signal source is at the left boundary. (**C**) Navigation success rate for cells using naive gradient tracking, receptor redistribution (Equation 5), Local Excitation Global Inhibition (LEGI), and minimal Bayes filter (Equation 4).

Cells simulated with receptor redistribution rapidly localize to the ligand source in patchy gradients, effectively avoiding signal traps (Figure 3B). We evaluate navigation efficiency by comparing simulations of four strategies: gradient tracking, receptor redistribution, bayes filtering, and Local Excitation-Global Inhibition (LEGI) (*16*). Gradient-tracking cells strictly follow the direction of local ligand gradient and become trapped in local peaks (Figure 3B), failing to cross the 60 µm gradient even after three hours. LEGI, which selectively amplifies spatial signals to track shallow gradients (*29*), also fails under these conditions. In contrast, receptor redistribution (and Bayes filtering) enable cells to consistently reach the source (Figure 3C), overcoming local traps by following the direction of maximal receptor activity (or belief) at each step, either through the bayesian update rule (Equation 4) or its implementation via receptor dynamics (Equation 5). All simulations used physiologically plausible parameters (*16*).

### Cellular implementation extends beyond standard Bayes filtering

A distinctive feature of cellular Bayes filtering (Equation 5), distinct from standard robotics implementations (Equation 1), is the coupling between observed signals and the motion model (Figure 4A). When mapping Bayesian filtering to receptor redistribution (Figure 1C), we found that membrane diffusivity of receptors corresponds to the variance of the motion model. In cells, receptor activation via ligand binding reduces receptor diffusivity through multiple mechanisms (Figure 4A), effectively lowering the variance parameter of the motion model as signal strength increases. This coupling extends the cell’s implementation from a standard filter to an adaptive one.

**Figure 4:**
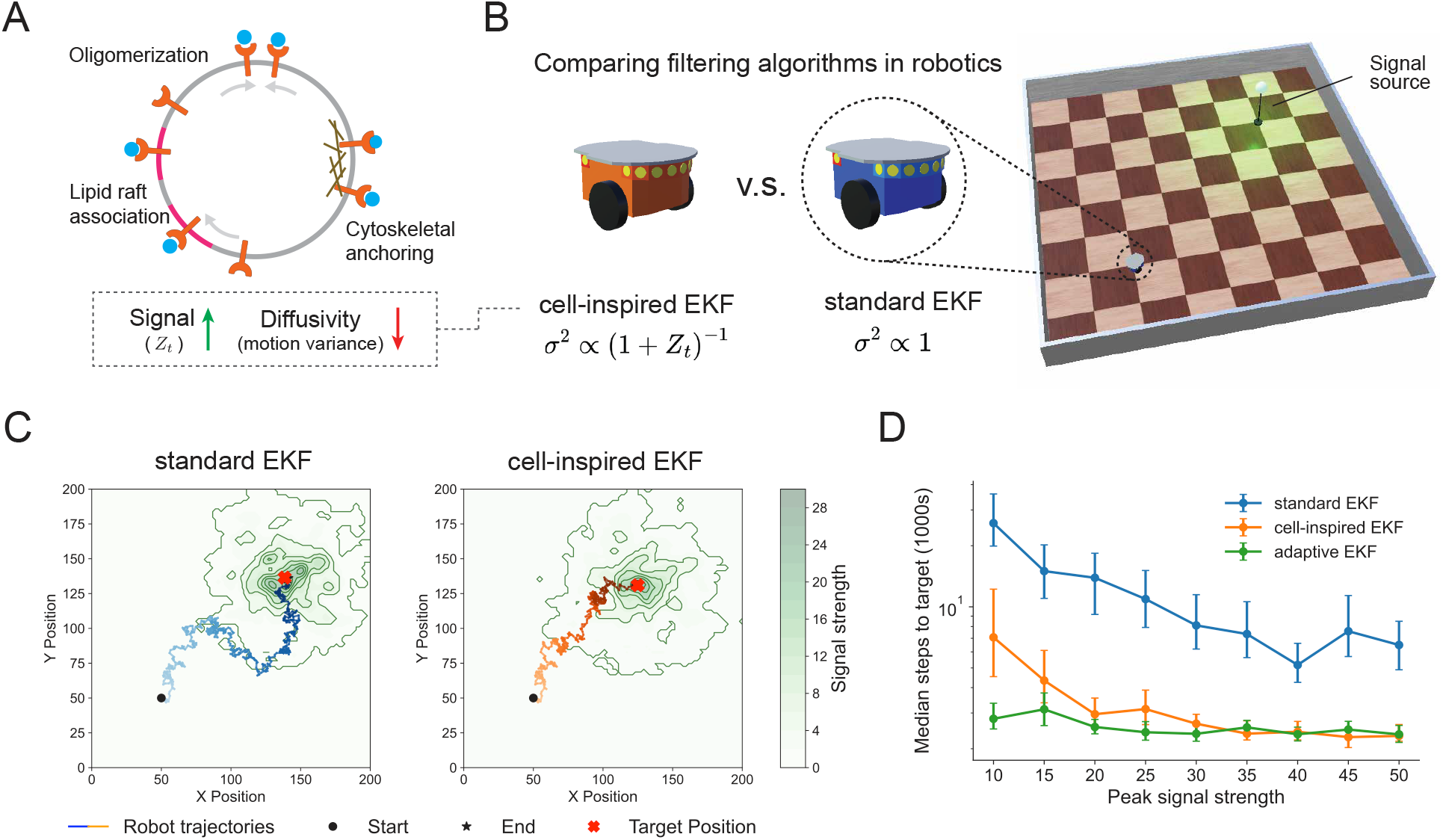
Cell-inspired, signal-coupled motion model for robotic navigation. (**A**) Multiple cellular mechanisms couple receptor activity (signal) with receptor diffusivity (motion model variance). (**B**) Image of simulated arena with robot and signal source. (**C**) Simulated trajectories of robots solving a target navigation task (*16*) using either standard Extended Kalman Filtering or coupled EKF which includes a signal-coupled motion model (Equation 6). (**D**) Navigation efficiency of three versions of EKF across environments with different peak signal strength; error bar represents SEM.

This signal-variance coupling improves source-seeking performance in environments with fluctuating noise. In robot simulations (Figure 4B), a standard Extended Kalman filtering (EKF) maintains a fixed motion model variance equal to the true motion variance,

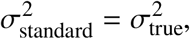

whereas our “cell-inspired EKF” adjusts variance according to the observed signal *Z*_*t*_,

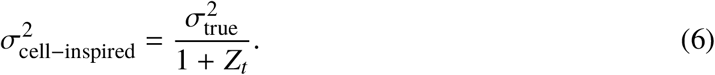

Incorporating this coupling reduced mean time-to-target 2–4 folds across a wide range of noise levels (Figure 4C, D).

We can understand the benefit of this receptorĐsignal coupling in terms of filter gain, which quantifies the degree to which new observations influence the belief (posterior) about the systemÕs state. For Poisson-distributed signals, stronger signal implies larger variability. In regimes of strong signal, therefore, cells and robots reduce their reliance on instantaneous observations by lowering the filter gain – precisely the effect achieved by decreasing motion model variance (*16*). It follows that an Adaptive EKF, which must continuously adjust the covariance term in the measurement likelihood to track changing noise statistics, achieves optimal performance. Remarkably, the cell-inspired EKF achieves nearly identical performance without the additional computation required to estimate measurement covariance (Figure 4D).

In conclusion, coupling receptor diffusivity to ligand engagement extends cellular Bayes filtering from a standard to an adaptive filter, enabling robust inference under fluctuating noise without the need for explicit noise estimation.

### Bayesian formalism predicts cell constraints

The mapping between the Bayesian formulation and the receptor PDE model predicts a coupling between cell speed and receptor dynamics.

In standard robotic implementations of Bayes filtering, the variance parameter *σ* of the motion model typically increases with robot speed (Equation 2). This relationship arises because faster motion accumulates greater positional uncertainty, as even small directional errors translate into larger spatial deviations over longer trajectories. Given that our mapping connects the motion model’s variance parameter *σ* to the receptor diffusivity *D*_*m*_ (Figure 1C), we predict that optimal receptor diffusivity should scale with cell speed. Indeed, our simulations confirm this prediction (Figure 5A), demonstrating that higher cell speeds require greater receptor diffusivity (Figure 5B). Intuitively, a fast-moving cell encounters new environmental signals more frequently and must thus rapidly revise its priors. Faster receptor diffusivity enables these rapid updates. Note Figure 5A also suggests that a fixed, low receptor diffusivity can still support efficient migration, despite being suboptimal.

**Figure 5:**
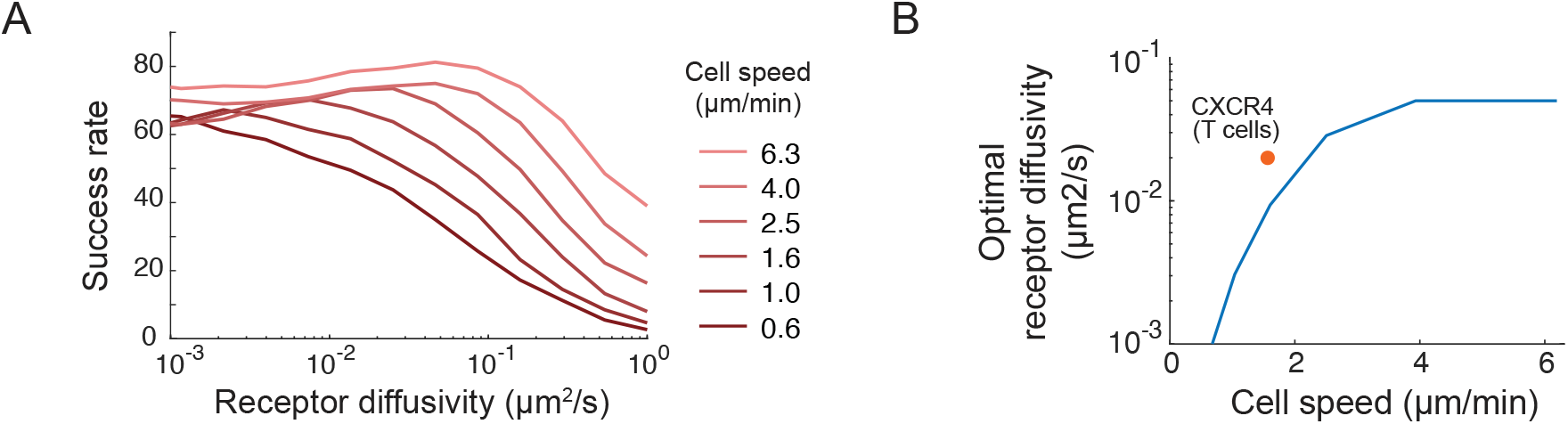
Bayesian formalism predicts optimal receptor diffusivity. (**A**) Success rate for cells simulated with different migration speed and receptor diffusivity. (**B**) Optimal receptor diffusivity at various cell speed, dot showing empirical data for CXCR4 in Jurkat T cells (*30, 31*).

## Discussion

In this work, we investigated the problem of source seeking by cells in complex tissue environments and established a direct mapping between receptor dynamics and Bayesian filtering, a widely used algorithm in robotic source seeking. This mapping shows that receptor redistribution can enable cells to efficiently navigate interstitial gradients, overcoming localized signal patches. This mapping also reveals a unique feature of the cellular Bayesian filter not present in standard Bayes filtering: the coupling between the observed signal and the motion model, which effectively acts as an adaptive Bayes filter enabling cells to adapt to fluctuating noise statistics.

Alternative implementations of Bayesian filtering in cells need not require receptor redistribution. The evolving belief distribution can instead be stored in the spatial distribution of signaling molecules that are recruited to the inner leaflet of the plasma membrane when receptors become active. In this scheme, the observed signal, *C*, driving the filter are receptor activation events, not extracellular ligand counts. After activation, many receptors (e.g., GPCRs) recruit cytosolic effectors, such as heterotrimeric G-proteins, adaptors, lipid kinases, to the membrane. Positive feedback loops (e.g. PI3K-Rac-F-action (*32*)) amplify and stabilize these effectors, effectively integrating the spatial distribution of recent receptor activity into the distribution of membrane-bound effectors. Because most membrane-bound effectors diffuse laterally, these molecules also naturally perform the prediction step involving the motion model, propagating the belief without requiring receptor relocation. Our Bayesian framework therefore applies to any membrane-associated species that (i) is produced or recruited in proportion to receptor activity and (ii) diffuses laterally, expanding the biochemical strategies cells might use to integrate noisy environmental cues.

Our work opens up new experimental directions. Future studies can leverage protein micropatterning to construct in vitro mimics of complex ligand landscapes observed in tissue, enabling simultaneous visualization of ligand distribution and the dynamics of surface receptors in migrating cells. These tools can be used to investigate how receptors in different cell types with different redistribution mechanisms (e.g., actin-vs. microtubule-based transport) differ in their responses to spatial signal structure.

Our work connects to a broader framework proposed by neuroscientist David Marr. Marr proposed that understanding an information-processing system requires analyzing it at three levels (*33*): the computational problem it solves, the algorithm it uses, and the physical implementation of that algorithm. This framework guides our analysis: starting from a navigation problem, we identify a Bayesian algorithm that solves it, and then show how mechanisms of receptor redistribution, as observed in cells, can implement this algorithm. Additional algorithmic strategies for decision making in complex environments may emerge from careful analysis of biological systems.

## Methods

### Derivation of equivalence between receptor PDE and Bayes filtering

The original discrete update equation is given as:

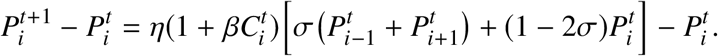

Rewriting and expanding:

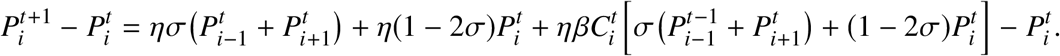

Grouping terms:

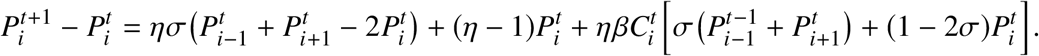

Replacing *P*_*i*_ with *R*_*i*_ and dividing by Δ*t*, we get:

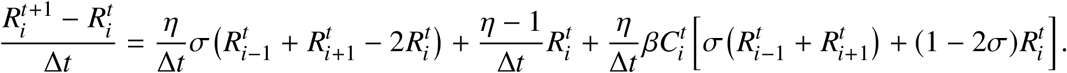

#### (a) Diffusion Term Mapping

The first term:

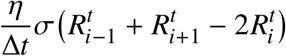

maps to diffusion. Given 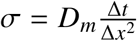 and *η* ≈ 1 for small Δ*t*, we simplify:

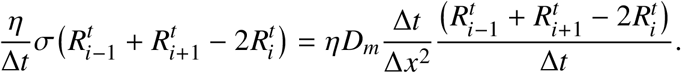

Cancelling Δ*t*:

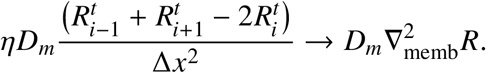

#### (b)Decay Term Mapping

The second term:

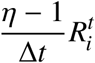

maps to receptor endocytosis. Given 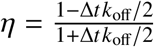:

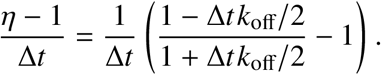

Simplifying:

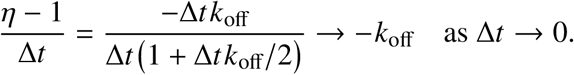

Thus:

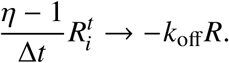

#### (c) Binding Term Mapping

The last term:

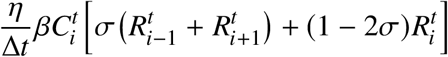

maps to receptor membrane transport. Given 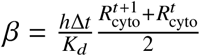 and 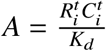, we get:

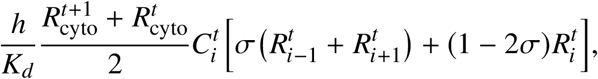

where *η* → 1 as Δ*t* → 0. Now, consider Δ*x* → 0, we can simplify this expression further

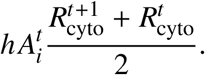

where 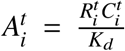. This is the semi-implicit treatment of *hAR*_cyto_ using the Crank-Nicolson method.

Finally, combining the terms, the final continuous-time PDE is:

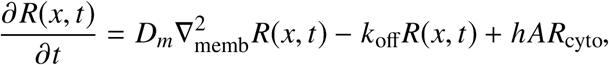

where 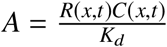.

### Fitting measurement likelihood to simulated tissue images

We obtain a measurement likelihood for the cellular bayesian filter by fitting to simulated images of interstitial gradients. The measurement likelihood is defined as *p*(*C*_*t*_ |*θ*_*t*_), which represents the likelihood of the cell observing ligand profile *C*_*t*_ ∈ ℤ^*N*^ around itself given a source direction *θ*_*t*_. For simplicity, we assume a local measurement likelihood where *P*(*C*_*t*_ |*θ*_*t*_) depends only on *C*_*t*_(*θ*_*t*_), the ligand level at direction *θ*_*t*_.

To evaluate the relationship between ligand concentration observations and localization likelihoods, we first sampled ligand concentration profiles from a simulated interstitial gradient (in the form of a ligand concentration field). Specifically, we defined a lattice of coordinates spanning the field and sampled ligand concentrations within a circular region of radius 10 µm, which approximates the region a cell would observe at different positions. For each sampled location, we extracted ligand concentrations over the boundary of the circular region. This produced a set of Poisson-distributed ligand count profiles *C*_*t*_ (*θ*_*t*_), which were used to represent signals observed by single cells.

Given these sampled ligand profiles, the probability *P*(*C*_*t*_ |*θ*_*t*_) of observing a particular ligand count at a given location was computed by marginalizing over all possible localization scenarios using shifted versions of the ligand field. These probabilities were then normalized across locations and converted to log-probabilities, log *P*(*C*_*t*_ |*θ*_*t*_). To analyze the statistical relationship between ligand concentration and localization probability, we fitted a linear regression model to predict log *P*(*C*_*t*_ |*θ*_*t*_) as a function of *C*_*t*_ (*θ*_*t*_) (Fig. S1) to obtain the following relationship:

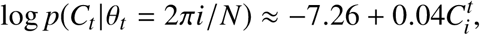

with a p-value of 1*e*^−3^, suggesting that the slope is significantly different from zero. Furthermore, since the coefficient of 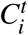 is small, we can perform a Taylor approximation of the exponential functio near zero to obtain:

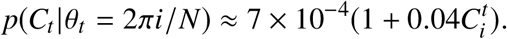

### Institial gradient simulation

We follow mathematical models of ligand distribution in tissue outlined in (*34, 35*), simulating a tissue environment using a PDE model that incorporates four transport mechanisms: (1) free diffusion, (2) ECM binding/unbinding, (3) fluid advection, (4) degradation. The spatial domain is a rectangle of size 300 µm × 900µm. We model ligands being supplied through fluid flows from the left boundary of the domain, and penetrate the interstitial space between immobilized cells. Soluble ligands are then transported by diffusion and fluid flow, and become immobilized upon binding to an extracellular matrix (ECM) made up of networks of interconnected fibers containing ligand binding sites. We explicitly represent both ECM-bound (*c*_*b*_) and soluble forms of the ligand (*c*_*s*_), so that the the total ligand concentration *c*(*x, t*) at position *x* and time *t* is equal to,

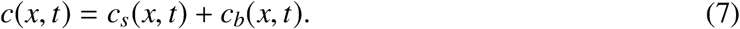

Mathematically, we can describe the dynamics of the soluble fraction *c*_*s*_(*x, t*) as follows,

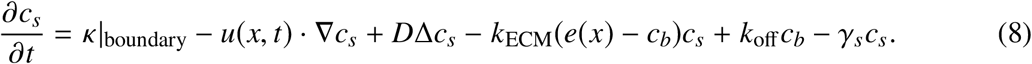

1. The first term, *κ*, represents production/release of molecule at the left boundary.
2. The second term represents fluid transport, where *u*(*x, t*) is the velocity field of the interstitial fluid with input flow speed *u*^in^ at the left boundary. We impose zero-velocity condition on the top and bottom boundary.
3. The third term represents diffusion with *D*_*m*_ as the ligand diffusion coefficient.
4. The fourth and fifth term represents ECM binding and unbinding, respectively. The concentration of ECM binding site *e*(*x*) at position *x* is generated using a minimal model of ECM protein distribution (see paragraph on “Generating ECM fiber network”). Binding occur with rate proportional to *e*(*x*) − *c*_*b*_(*x, t*), the level of available ECM binding site.
5. The last term represents enzymatic degradation of ligand.

The dynamics of ECM-bound fraction *c*_*b*_(*x, t*) is much simpler, involving ECM binding, ECM unbinding, and enzymatic decay.

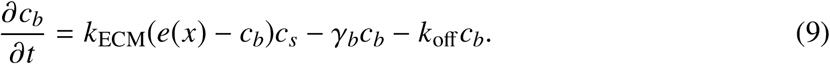

To generate a ligand concentration field *c*, we take *κ* to be non-zero for a brief period of time, representing a bolus of ligand released. Then, we simulate the combined dynamics of bound and soluble fractions for sufficiently long until the ligand distribution *c*(*x, t*) is relatively stable. In practice, we observe that *c* ≈ *c*_*b*_ after a sufficiently long period of time, since the soluble fraction quickly become insignificant due to fluid flow. The resulting concentration field represents an interstitial gradient. The average concentration is set by setting the release rate *κ* such that the concentration of the soluble fraction *c*_*s*_ matches measured chemokine concentration found in interstitial fluids (1-10 pM) (*36, 37*).

#### Generating ECM fiber network

To generate a distribution of ECM binding sites *e*(*x*) Equation (9), we use a minimal computation model of fiber network (*38–40*). The model generates ECM fibers represented by line segments, which could represent fibronectin, collagen, laminin, or other fibrous matrix components. To position each fiber, one end of each segment is randomly positioned following a uniform distribution within the domain. The other end’s position is determined by picking an angle, uniformly from [0, 2*π*), and length sampled from a normal distribution with mean 75µ*m* and standard deviation of 5µ*m* (as measured for collagen by Friedl et al (*41*)). In total, 4050 fibers were placed in the domain. For the PDE simulation, the generated network is discretized by counting the number of fibrous proteins around each node in the simulation lattice. The density of fiber within each node is then converted to a concentration value representing the level of ECM binding sites, resulting in an average concentration of ECM binding site of 520 nM.

### Numerical simulation of receptor redistribution dynamic

Receptor *r*(*x, t*) is modeled by considering three redistribution mechanisms: (1) lateral diffusion of *r* along the plasma membrane 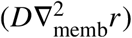, (2) endocytosis of *r* along the plasma membrane (*k*_off_ *r*), (3) incorporation of cytoplasmic pool of receptors, *R*_cyto_, to the membrane at rate proportional to local receptor activity (*h AR*_cyto_). *A*(*x, t*) is a random variable that denotes receptor activity along the cell membrane, and is a function of local receptor number. Then, the equation describing the distribution of *r* across the cell membrane can be expressed mathematically as,

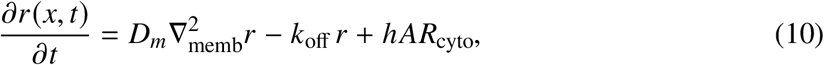

where the total number of receptors *r*_tot_ =∫_memb_ *r* + *R*_cyto_ is fixed.

We simulate receptor distribution by treating the cell membrane as a 1D space and the cytosol as a single, homogeneous compartment. This simplification allows us to simulate our PDE using the Crank-Nicolson method in one spatial dimension. Given space and time units Δ*x* and Δ*t*, respectively, the Crank-Nicolson method with 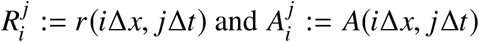 is given by the difference scheme

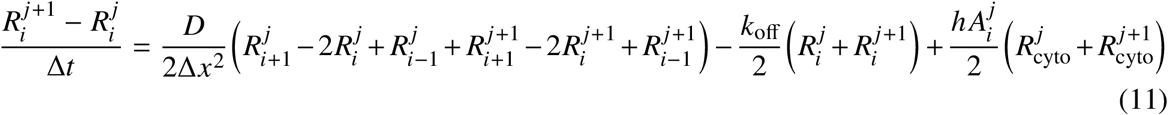

where, *i* = 1, 2, 3, ∼*m*, representing *m* discrete membrane compartments and 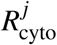 represents the additional cytosol compartment. Since the membrane is represented by a circle, we have the following pair of conditions,

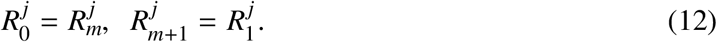

Lastly, total receptor number across all compartments is conserved,

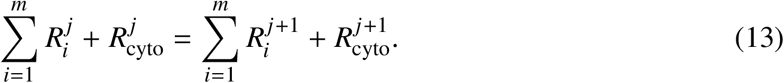

Now, we can combined Equation (11)-(13) and rewrite everything in vector form. First, let

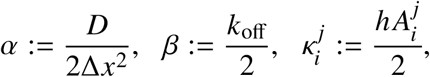

and rewrite Equation (11) as,

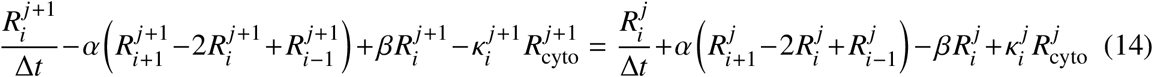

and define *U*^*j*^ to be the (*m* + 1)-dimensional vector with components 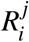 for *i* = 1, 2, 3, ∼*m* and 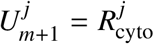. The difference scheme is given in the vector form

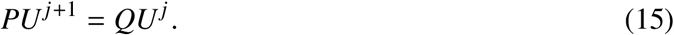

where,

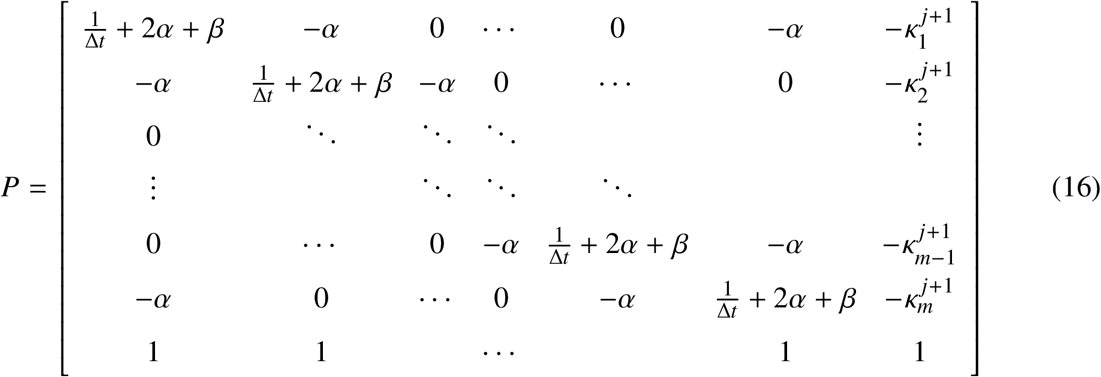

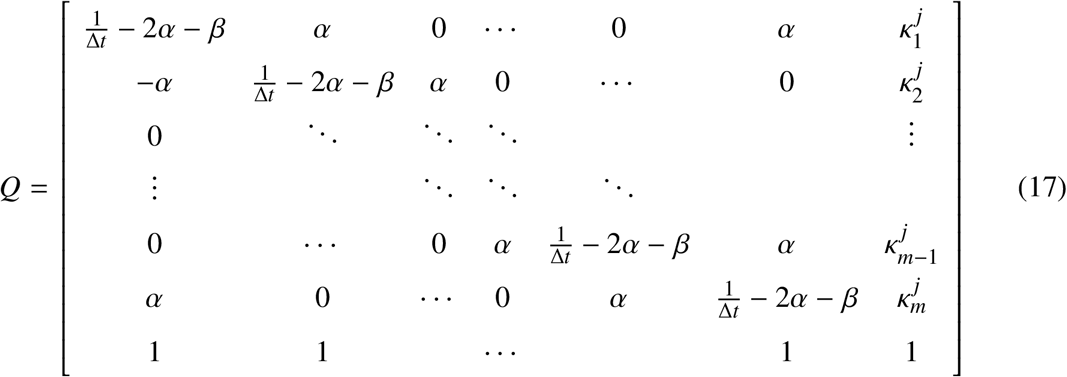

Because *A* is invertible, the Crank-Nicolson scheme reduces to the iterative process

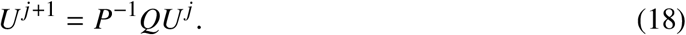

Furthermore, we model receptor activation *A*_*i*_ as follows (*42*),

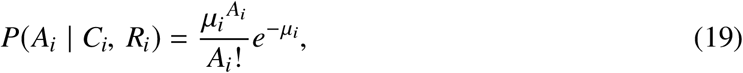

where 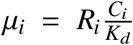. The bracket term represents the probability of activation for a receptor experiencing *C*_*i*_ ligands. *K*_*d*_ is the equilibrium dissociation constant. In other words, the number of active receptors *A*_*i*_ given ligand count *C*_*i*_ is a Poisson random variable with mean µ_*i*_. The entire evolution of *r* can be solved where at each time step, we update receptor activity 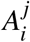 across all membrane position *i* according to the random process described by Equation (19), followed by solving Equation (18) for *U*^*j*+1^.

We set the value of the feedback constant *h* using empirical measurements from (*43*). In Figure 3M of Marco et al., the authors report a quartile box plot showing estimated values for a parameter they call h (which we will refer to as 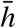), with a mean estimate of around 1.6 × 10^−3^/*s*. Note 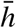 is equivalent in meaning as our *hA*_*i*_. However, since *hA*_*i*_ will be different across different membrane bins and across time, we simulate the feedback scheme for a cell in a given environment and set the value of h such that the mean rate ⟨*hA*_*i*_⟩ (averaged across membrane and time) is approximately equal to the mean estimate of 1.6 × 10^−3^ reported by Marco et al.. The value 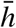 reported by Marco et al. corresponds specifically to the transport rate of the Cdc42 to the membrane. The parameter value was obtained by analyzing fluorescence recovery of GFP-Cdc42 in membrane regions bleached with a laser pulse. Although the measured value corresponds to Cdc42, it has been used to model the effective recruitment rate for receptors shown to undergo polarized exocytosis, showing good agreement with empirical data (*44*). Similar values around 1 − 2 × 10^−3^/*s* have been measured for the recycling rate of a wide range of GPCRs (*45–48*).

### Numerical simulation of Local Excitation and Global inhibition

We simulate a Local Excitation, Global Inhibition Biased Excitable Network (LEGI-BEN) as described by Shi et al. (*49*). This model implements an excitable network module based on an activator-inhibitor system, where stochastic fluctuations initiate activity. The dynamics are governed by the following equations:

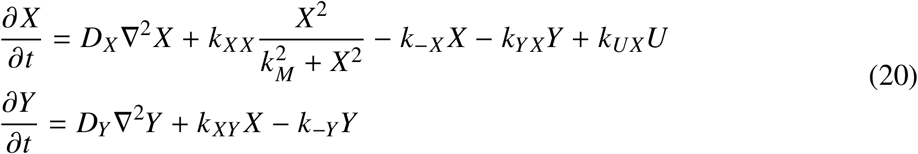

Here, U serves as the input to the excitable system and includes contributions from three components: basal activation (B), stochastic fluctuations (N), and the response regulator (R) from the LEGI module, as shown below:

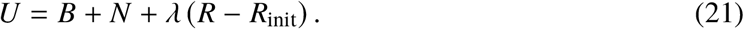

The LEGI module processes the stimulus (S) to drive the response regulator (R), which biases the activity of the excitable network. Its dynamics are described by the following system:

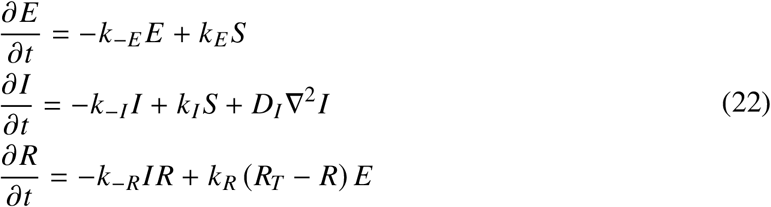

In this framework, the activity of Y determines the direction of cell migration, with cells moving toward the region of maximal Y activity on their surface.

All simulations and the model implementation were carried out in MATLAB (*50*), closely following the protocol in (*49*). The signaling partial differential equations (PDEs) were solved on a one-dimensional representation of the cell boundary, using periodic boundary conditions. The spatial domain was discretized into 360 points, and spatial diffusion terms were approximated with central finite differences. This discretization transforms the PDEs into ordinary differential equations (ODEs), which were solved using the SDE toolbox in MATLAB. Simulations were run with a time step of 0.025 seconds.

### Simulation of cell navigation

#### Navigation algorithm

At *t* = 0, initialize a cell at position *p*_0_ ∈ Ω ⊂ ℝ^2^. At each subsequent time step *t* = *t* + Δ*t* with the cell at position *p*_*t*_ ∈ Ω:

1. Compute mean ligand profile ***c*** ∈ ℝ^*m*^ at the cell’s current position.
2. Independently sample *n* ligand profiles 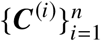 where each element *C*_*j*_ is distributed as a Poisson random variable with mean equal to *c*_*j*_.
3. For each ligand profile ***C***^(*i*)^ sampled, sample a corresponding receptor activity profiles ***A***^(*i*)^,

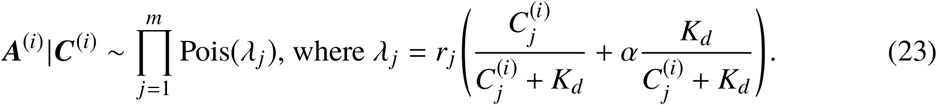
4. Update belief distribution if relevant.
5. Select the next direction 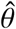 as the direction of maximal receptor activity or belief probability in the case of Bayes filtering.
6. Set new cell position 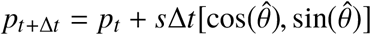, with speed *s* = 1µm/min, Δ*t* = 1*s*.
7. Repeat from step 1.

### Robot navigation simulation

We implemented an Extended Kalman Filter (EKF) for target navigation where a robot localizes and moves toward a target using noisy signal measurements. A Kalman filter is a special type of Bayes filter where both the motion and measurement noise are Gaussian and are more amenable to analytical treatments. Furthermore, an Extended Kalman Filter is an extension of Kalman filter to the case of nonlinear process and/or measurement model, using linearized model via Taylor series expansion.

### Environment setup

The environment is a 200 × 200 arena, with the robot starting at (50, 50) and the target starting at (120, 120). The target emits an signal which decays exponentially from the target. Specifically, the expected signal level at position *x* is

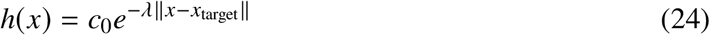

where *c*_0_ is the maximum signal strength, *λ* is the decay exponent. A robot navigating in this environment observes signal *Z*_*t*_, which is a Poisson random variable with mean *h*(*x*),

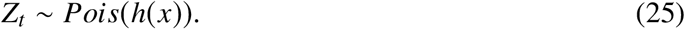

#### Target dynamics

The target undergoes a 2D random walk at each time step. The target position evolves according to:

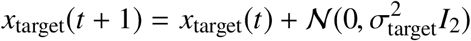

where *σ*_target_ is the target motion standard deviation and *I*_2_ is the 2 × 2 identity matrix. The target position is constrained to remain within the grid boundaries [0, 200) × [0, 200).

#### Robot action

The robot maintains a belief state *b*_*t*_(*x*) representing a probability distribution over possible target locations which gets updated per unit time. At each iteration, the robot updates its belief by observing a signal at its own grid position *r*_*t*_ in the arena, and moves in the direction of the maximum likelihood target location, subject to actuator noise represented by Gaussian noise.

#### Belief update

The robots’ belief *b*_*t*_(*x*) is updated in two stages: prediction and update. The prediction step incorporates target motion uncertainty:

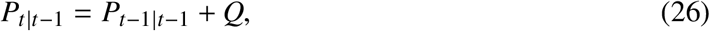

where P is the covariance of the belief distribution at time *t* − 1, and *Q* is the covariance of the motion noise. For standard EKF, we set *Q* to be the true motion covariance 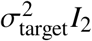. For the coupled strategy, the process variance *σ*^2^ varies with signal *Z*_*t*_:

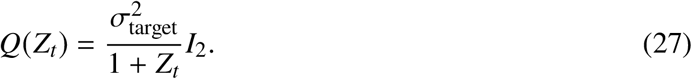

The update step consists of the following:

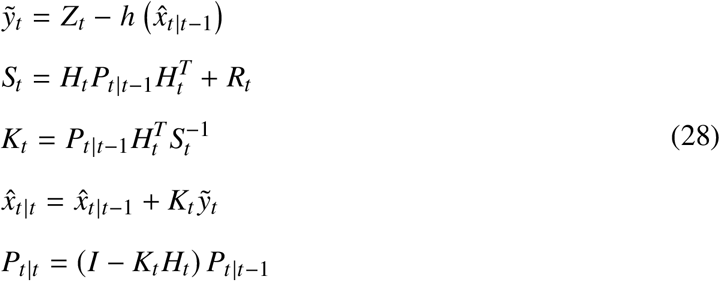

where 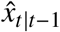is the previous state estimate, 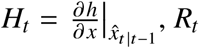 is the measurement noise which we take to be the true signal noise at the robot’s initial position.

#### Adaptive filtering

As an extension, we compare both strategies to adaptive EKF which dynamically adjust measurement noise parameters based on innovation statistics. The measurement variance *R*_*t*_ is updated using exponential smoothing:

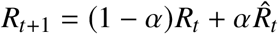

where *α* is the adaptation rate and 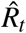 is the empirical measurement noise estimate computed from the innovation sequence 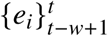 over a sliding window of size w:

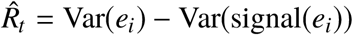

This allows the filter to adapt to changing observation statistics during navigation.

#### Performance Evaluation

We compared different strategies through multiple simulation runs across environments with different peak signal strength. A run was considered successful if the robot reached within three unit distance of the target. All strategies were evaluated using identical random seeds.

#### Understanding coupled filtering strategy based on Kalman gain

We can understand the effectiveness of the adaptive strategy by studying the gain of a 1-D Kalman filter. The Kalman gain (*K*_*t*_) controls how much weight is given to a new measurement relative to the current belief, defined as:

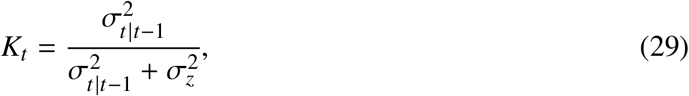

where 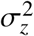is the variance of the measurement noise and 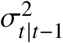 is the variance of the previous estimate. The variance term after motion update 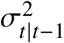is the sum of the posterior variance from the previous step and the variance of the motion model:

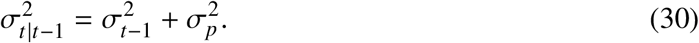

Combining Equation (29) and Equation (30), we rewrite the Kalman gain:

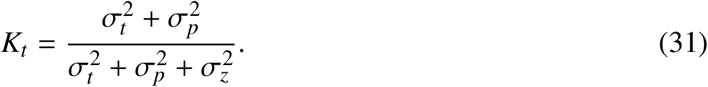

The Kalman gain takes a value between 0 and 1, where a larger gain means that new measurements are weighted more heavily relative to the prior belief.

With standard Kalman filtering, 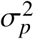 and 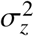are both assumed to be constant. Thus, given some prior belief distribution, the Kalman gain eventually converges to a fixed limit as obtained from the algebraic Ricatti equation. For our simulated environment with Poisson noise (Equation 25), however, signal noise scales with signal strength which varies throughout the environment (Equation 24). Therefore, *K*_*t*_ should be dynamically tuned as the signal noise changes. Specifically as 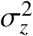increases, the Kalman gain should decrease, representing the fact that the signals are becoming noisier.

Recall 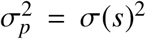 decreases as the signal *s* becomes stronger according to Equation (27). Thus this coupling causes the Kalman gain to decrease as signal increases, which is precisely what one should do for Poisson noise. An alternative strategy to deal with fluctuating measurement noise is to continuously estimate and revise the signal noise 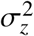 used by the filtering algorithm, which is what adaptive kalman filter aims to do. Indeed, we observe that the cell-inspired coupling strategy matches the performance of adaptive filtering without the additional computation required for continuous noise estimation.

## Acknowledgments

We would like to thank Pablo Iglesias for providing the LEGI-BEN simulation code.

## Author contributions

Z. J. W. conceived the idea and performed all simulation and mathematical analysis. Z. J. W. and M. T. wrote the paper. Both Z. J. W. and M.T. provided funding for the project.

## Data and materials availability

The code used in this study is publicly available on GitHub at <monospace>cellethology/bayesian-cell</monospace>. This repository contains all scripts and instructions necessary to reproduce the primary results presented in the paper.

## Extended Data

Tables S1 to S3

Figure S1

## Extended Data

**Figure S1:**
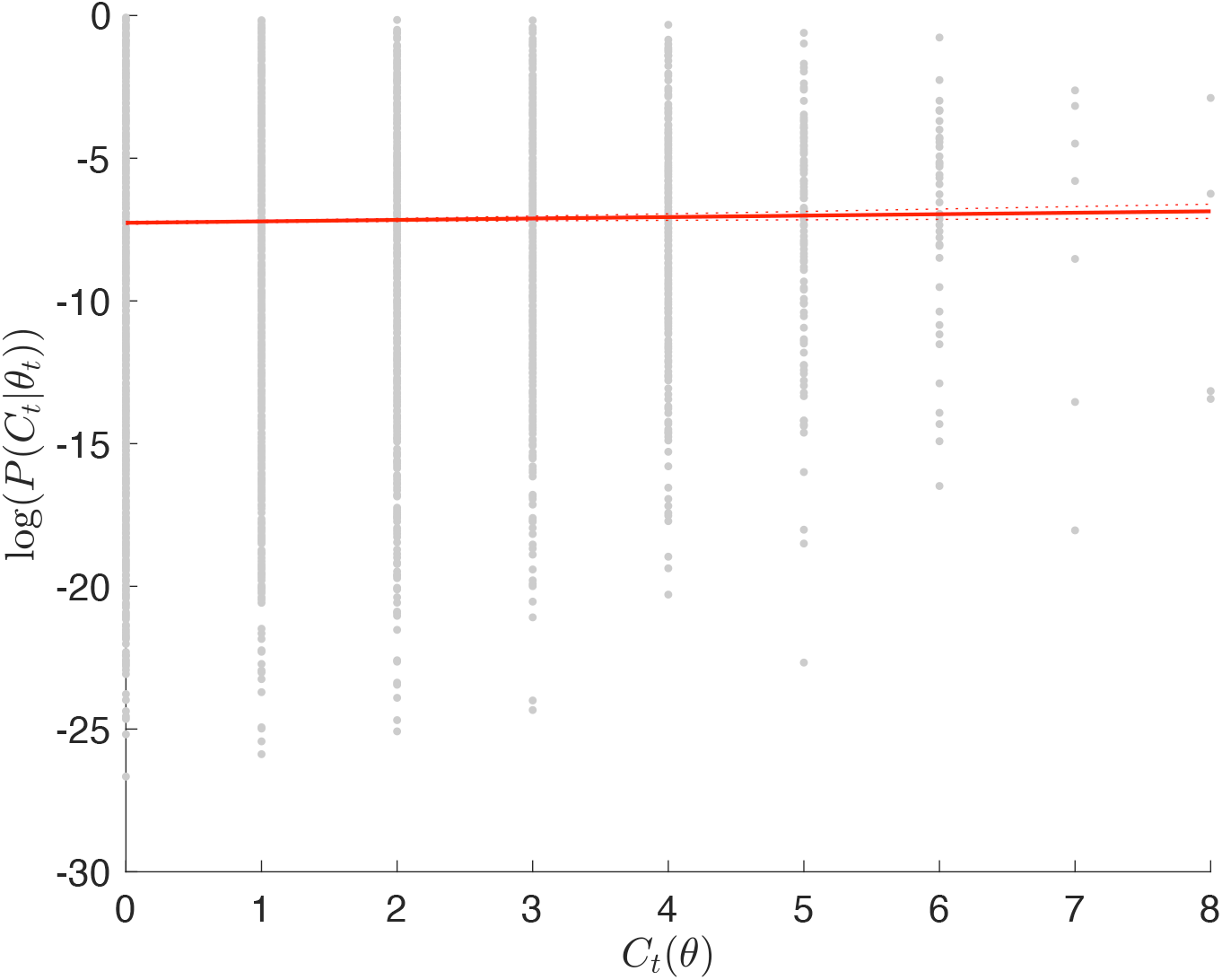
Fitting a local measurement likelihood model for navigating interstitial gradients. Each point of scatter plot represents a ligand profile as observed by a 10 µm cell in a simulated interstitial gradient where the signal originates from a direction *θ*_*t*_. Red solid line is the fitted linear regression line, dashed lines indicate the 95% confidence interval. Fitted linear model with p-value = 0.001.

**Table S1:** Parameters used for modeling tissue gradient. Including parameters for fluid simulation (*34*) and ECM network (*39*).

| Parameter | Symbol | Value |
| --- | --- | --- |
| Domain size | — | $300\ \mu\text{m} \times 900\ \mu\text{m}$ |
| Diffusion coefficient | $D$ | $45\ \mu\text{m}^2\text{s}^{-1}$ |
| Fluid viscosity | $\mu$ | $2.5\ \mu\text{g}\mu\text{m}^{-1}\text{s}^{-1}$ |
| Spatial discretization | $\Delta x$ | $2\ \mu\text{m}$ |
| Time step | $\Delta t$ | $0.0178\ \text{s}$ |
| Interstitial fluid input flow | $u^{\text{in}}$ | $1\ \mu\text{m}\ \text{s}^{-1}$ |
| Average [ECM binding site] | — | $520\ \text{nM}$ (51) |
| ECM binding rate | $k_{\text{ECM}}$ | $9.3 \times 10^{-5}\ \text{nM}^{-1}\text{s}^{-1}$ (52) |
| ECM release rate | $k_{\text{off}}$ | $1.2 \times 10^{-4}\ \text{s}^{-1}$ (52) |
| Production/release rate | $\kappa$ | $7\ \text{nM}\ \text{s}^{-1}$ |
| Soluble ligand degradation | $\gamma_s$ | $1 \times 10^{-3}\ \text{s}^{-1}$ (51) |
| Bound ligand degradation | $\gamma_b$ | $1 \times 10^{-5}\ \text{s}^{-1}$ (35) |
| Mean ECM fiber length | — | $75\ \mu\text{m}$ |
| Variance in fiber length | — | $5\ \mu\text{m}$ |
| Total number of ECM fibers | — | 4050 |

**Table S2:**
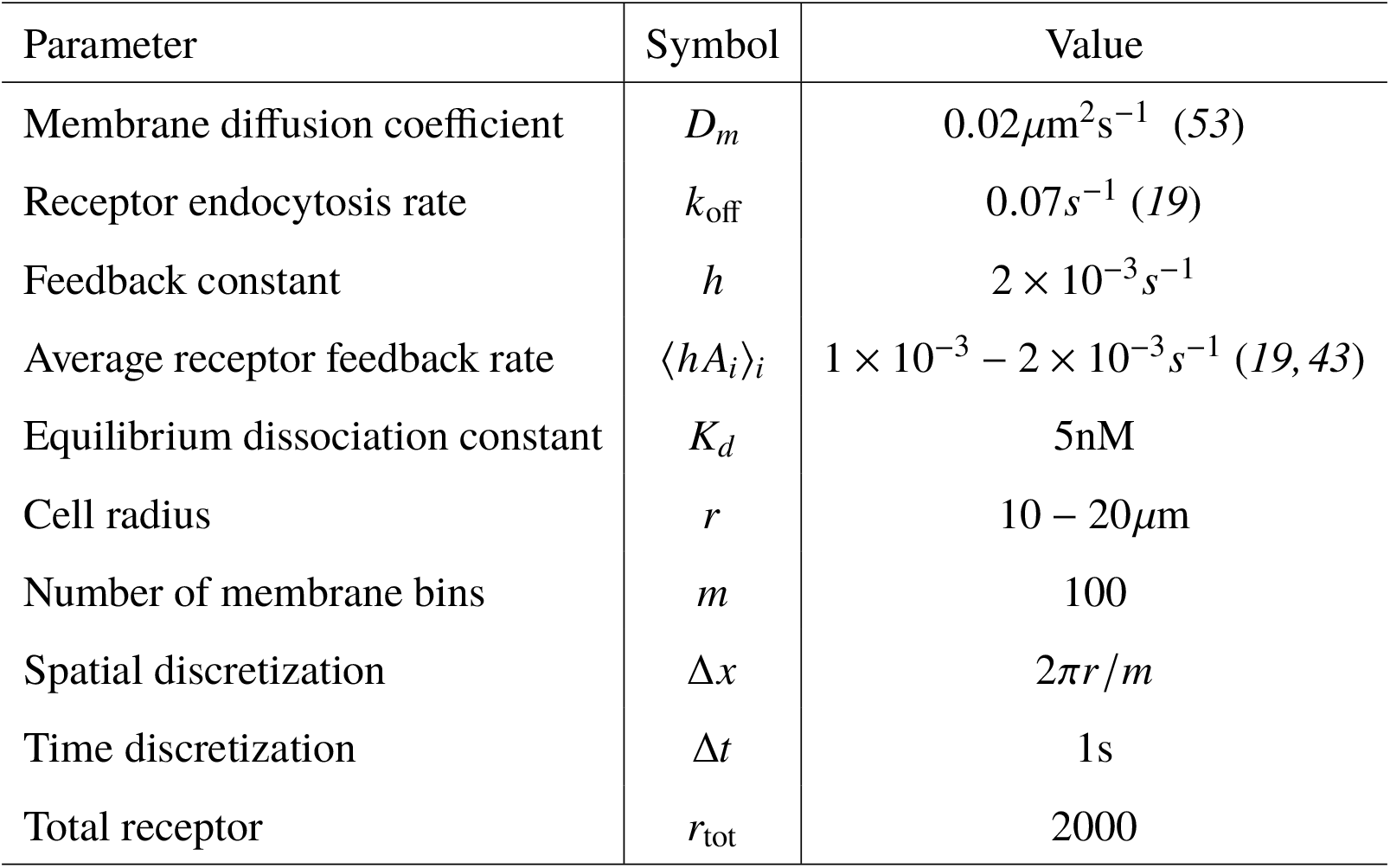
Parameter values for simulating receptor redistribution. The average rate of receptor incorporation, ⟨*h A*_*i*_⟩_*i*_, depends on receptor activity which changes as the cell moves in a heterogeneous environment. Thus, the range shown represents the time-averaged value for a cell moving through simulated tissue environments. The value of *h* was chosen to achieve a physiologically relevant range for ⟨*h A*_*i*_⟩_*i*_.

**Table S3:**
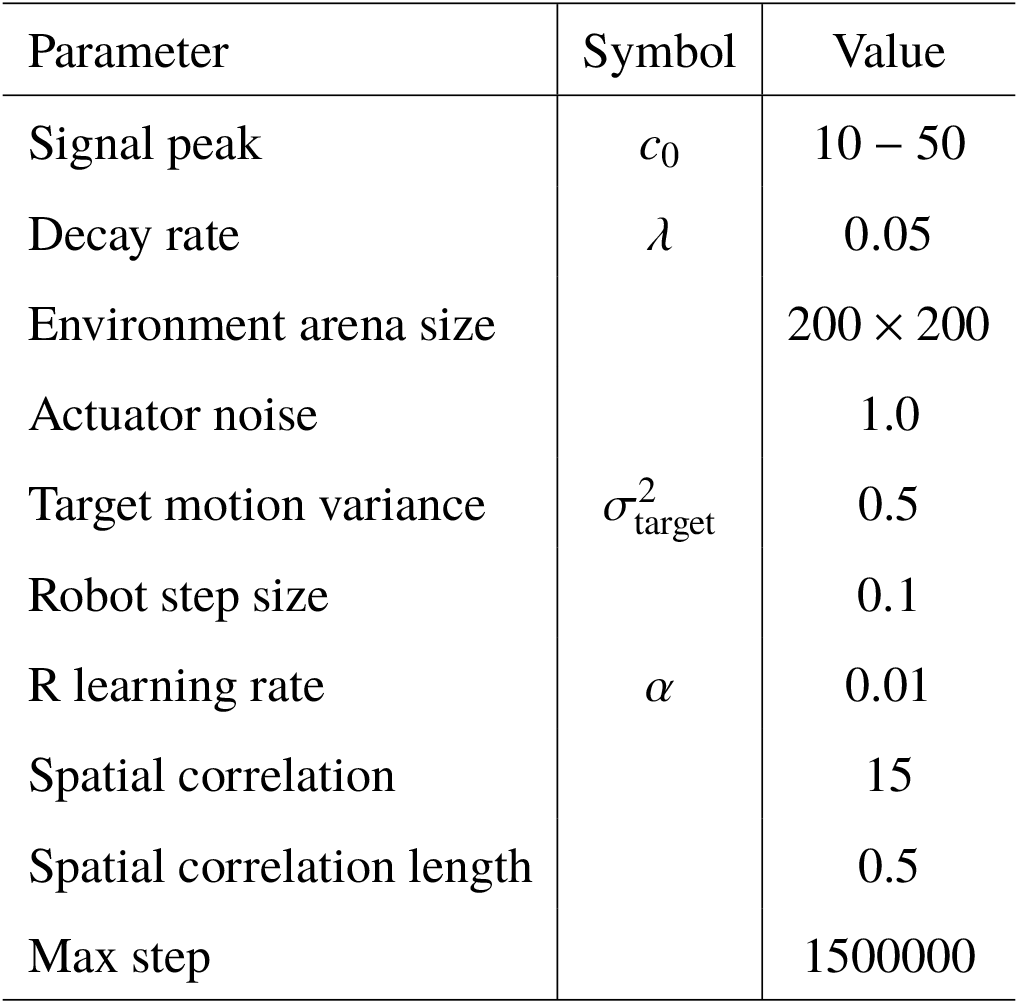
Parameter values for robotic simulation with extended kalman filter.

## References and Notes

1. A. Francis, S. Li, C. Griffths, J. Sienz, Gas source localization and mapping with mobile robots: A review. Journal of Field Robotics 39 (8), 1341–1373 (2022).

2. W. Naeem, R. Sutton, J. Chudley, Chemical plume tracing and odour source localisation by autonomous vehicles. The Journal of Navigation 60 (2), 173–190 (2007).

3. S. Thrun, W. Burgard, D. Fox, Probabilistic robotics (MIT Press, Cambridge, Mass.) (2005).

4. D. J. Fowell, M. Kim, The spatio-temporal control of effector T cell migration. Nature Reviews Immunology pp. 1–15 (2021).

5. E. Russo, et al., Intralymphatic CCL21 promotes tissue egress of dendritic cells through afferent lymphatic vessels. Cell reports 14 (7), 1723–1734 (2016).

6. A. C. von Philipsborn, et al., Growth cone navigation in substrate-bound ephrin gradients. Development 133 (13), 2487–2495 (2006).

7. M. Weber, et al., Interstitial dendritic cell guidance by haptotactic chemokine gradients. Science 339 (6117), 328–332 (2013).

8. Z. J. Wang, M. Thomson, Localizaion of signaling receptors maximizes cellular information acquisition in spatially structured natural environments. Cell Systems 13 (7), 530–546 (2022).

9. T. E. Sutherland, D. P. Dyer, J. E. Allen, The extracellular matrix and the immune system: A mutually dependent relationship. Science 379 (6633), eabp8964 (2023).

10. T. E. Kennedy, H. Wang, W. Marshall, M. Tessier-Lavigne, Axon guidance by diffusible chemoattractants: a gradient of netrin protein in the developing spinal cord. Journal of Neuro-science 26 (34), 8866–8874 (2006).

11. M. Sarris, et al., Inflammatory chemokines direct and restrict leukocyte migration within live tissues as glycan-bound gradients. Current Biology 22 (24), 2375–2382 (2012).

12. E. Donà, et al., Directional tissue migration through a self-generated chemokine gradient. Nature 503 (7475), 285–289 (2013).

13. C. Guirland, S. Suzuki, M. Kojima, B. Lu, J. Q. Zheng, Lipid rafts mediate chemotropic guidance of nerve growth cones. Neuron 42 (1), 51–62 (2004).

14. A. Pignata, et al., A spatiotemporal sequence of sensitization to slits and semaphorins orchestrates commissural axon navigation. Cell reports 29 (2), 347–362 (2019).

15. F. Belema-Bedada, S. Uchida, A. Martire, S. Kostin, T. Braun, Effcient homing of multipotent adult mesenchymal stem cells depends on FROUNT-mediated clustering of CCR2. Cell stem cell 2 (6), 566–575 (2008).

16. Materials and methods are available as supplementary material.

17. C. C. Winkle, et al., A novel Netrin-1–sensitive mechanism promotes local SNARE-mediated exocytosis during axon branching. Journal of Cell Biology 205 (2), 217–232 (2014).

18. C. Bouzigues, M. Morel, A. Triller, M. Dahan, Asymmetric redistribution of GABA receptors during GABA gradient sensing by nerve growth cones analyzed by single quantum dot imaging. Proceedings of the National Academy of Sciences 104 (27), 11251–11256 (2007).

19. A. W. McClure, et al., Role of polarized G protein signaling in tracking pheromone gradients. Developmental cell 35 (4), 471–482 (2015).

20. M. Shimonaka, et al., Rap1 translates chemokine signals to integrin activation, cell polarization, and motility across vascular endothelium under flow. The Journal of cell biology 161 (2), 417– 427 (2003).

21. T. Tojima, et al., Attractive axon guidance involves asymmetric membrane transport and exocytosis in the growth cone. Nature neuroscience 10 (1), 58–66 (2007).

22. P. Wan, et al., Guidance receptor promotes the asymmetric distribution of exocyst and recycling endosome during collective cell migration. Development 140 (23), 4797–4806 (2013).

23. J. Zeng, S. Feng, B. Wu, W. Guo, Polarized exocytosis. Cold Spring Harbor perspectives in biology 9 (12), a027870 (2017).

24. T. Cotrufo, et al., A signaling mechanism coupling netrin-1/deleted in colorectal cancer chemoattraction to SNARE-mediated exocytosis in axonal growth cones. Journal of Neuroscience 31 (41), 14463–14480 (2011).

25. G. Servant, O. D. Weiner, E. R. Neptune, J. W. Sedat, H. R. Bourne, Dynamics of a chemoattractant receptor in living neutrophils during chemotaxis. Molecular biology of the cell 10 (4), 1163–1178 (1999).

26. G. Servant, et al., Polarization of chemoattractant receptor signaling during neutrophil chemotaxis. Science 287 (5455), 1037–1040 (2000).

27. Z. Wang, et al., UNC-6 (netrin) stabilizes oscillatory clustering of the UNC-40 (DCC) receptor to orient polarity. Journal of Cell Biology 206 (5), 619–633 (2014).

28. Z. Wang, et al., Extracellular vesicles in fatty liver promote a metastatic tumor microenvironment. Cell metabolism 35 (7), 1209–1226 (2023).

29. Y. Xiong, C.-H. Huang, P. A. Iglesias, P. N. Devreotes, Cells navigate with a local-excitation, global-inhibition-biased excitable network. Proceedings of the National Academy of Sciences 107 (40), 17079–17086 (2010).

30. X. Yin, D. A. Knecht, M. A. Lynes, Metallothionein mediates leukocyte chemotaxis. BMC immunology 6 (1), 21 (2005).

31. E.M. García-Cuesta, et al., Allosteric modulation of the CXCR4: CXCL12 axis by targeting receptor nanoclustering via the TMV-TMVI domain. Elife 13, RP93968 (2024).

32. T. Inoue, T. Meyer, Synthetic activation of endogenous PI3K and Rac identifies an AND-gate switch for cell polarization and migration. PloS one 3 (8), e3068 (2008).

33. D. Marr, T. Poggio, From Understanding Computation to Understanding Neural Circuitry, Tech. rep., USA (1976).

34. K. A. Rejniak, et al., The role of tumor tissue architecture in treatment penetration and effcacy: an integrative study. Frontiers in oncology 3, 111 (2013).

35. F. Milde, M. Bergdorf, P. Koumoutsakos, A hybrid model for three-dimensional simulations of sprouting angiogenesis. Biophysical journal 95 (7), 3146–3160 (2008).

36. X. Wang, M. R. Lennartz, D. J. Loegering, J. A. Stenken, Multiplexed cytokine detection of interstitial fluid collected from polymeric hollow tube implantsâATa feasibility study. Cytokine 43 (1), 15–19 (2008).

37. K. E. Clark, et al., Multiplex cytokine analysis of dermal interstitial blister fluid defines local disease mechanisms in systemic sclerosis. Arthritis research & therapy 17 (1), 1–11 (2015).

38. D. Harjanto, M. H. Zaman, Modeling extracellular matrix reorganization in 3D environments. PLoS One 8 (1), e52509 (2013).

39. D. K. Schlüter, I. Ramis-Conde, M. A. Chaplain, Computational modeling of single-cell migration: the leading role of extracellular matrix fibers. Biophysical journal 103 (6), 1141– 1151 (2012).

40. B. Lee, et al., A three-dimensional computational model of collagen network mechanics. PloS one 9 (11), e111896 (2014).

41. P. Friedl, et al., Migration of highly aggressive MV3 melanoma cells in 3-dimensional collagen lattices results in local matrix reorganization and shedding of α2 and β1 integrins and CD44. Cancer research 57 (10), 2061–2070 (1997).

42. M. Ueda, Y. Sako, T. Tanaka, P. Devreotes, T. Yanagida, Single-molecule analysis of chemotactic signaling in Dictyostelium cells. Science 294 (5543), 864–867 (2001).

43. E. Marco, R. Wedlich-Soldner, R. Li, S. J. Altschuler, L. F. Wu, Endocytosis optimizes the dynamic localization of membrane proteins that regulate cortical polarity. Cell 129 (2), 411– 422 (2007).

44. B. Hegemann, et al., A cellular system for spatial signal decoding in chemical gradients. Developmental cell 35 (4), 458–470 (2015).

45. L. D. L. JJ, Receptors: Models for binding, traffcking, and signaling (1993).

46. S. Pippig, S. Andexinger, M. J. Lohse, Sequestration and recycling of beta 2-adrenergic receptors permit receptor resensitization. Molecular pharmacology 47 (4), 666–676 (1995).

47. J. A. Koenig, J. M. Edwardson, Intracellular traffcking of the muscarinic acetylcholine receptor: importance of subtype and cell type. Molecular pharmacology 49 (2), 351–359 (1996).

48. J. A. Koenig, J. M. Edwardson, Kinetic analysis of the traffcking of muscarinic acetylcholine receptors between the plasma membrane and intracellular compartments. Journal of Biological Chemistry 269 (25), 17174–17182 (1994).

49. C. Shi, C.-H. Huang, P. N. Devreotes, P. A. Iglesias, Interaction of motility, directional sensing, and polarity modules recreates the behaviors of chemotaxing cells. PLoS computational biology 9 (7), e1003122 (2013).

50. MATLAB, 23.2.0.2409890 (R2023b) (The MathWorks Inc., Natick, Massachusetts) (2023).

51. Y. Wang, D. J. Irvine, Convolution of chemoattractant secretion rate, source density, and receptor desensitization direct diverse migration patterns in leukocytes. Integrative Biology 5 (3), 481–494 (2013).

52. J. D. Shields, et al., Autologous chemotaxis as a mechanism of tumor cell homing to lymphatics via interstitial flow and autocrine CCR7 signaling. Cancer cell 11 (6), 526–538 (2007).

53. L. Martínez-Muñoz, et al., Separating actin-dependent chemokine receptor nanoclustering from dimerization indicates a role for clustering in CXCR4 signaling and function. Molecular cell 70 (1), 106–119 (2018).

